# Wavelet Based Whole Genome Doubling Aware Single Cell Copy Number Calling

**DOI:** 10.64898/2025.12.18.693686

**Authors:** Benjamin K. Wesley, Frank Wos, Soren Germer, Jade E.B. Carter, Silas Maniatis, Khanh Dinh, James S. Roche, Timothy R. Chu, Nicolas Robine, Rebecca Fitzgerald, John Lizhe Zhuang, Simon Tavaŕe, Karol Nowicki-Osuch

## Abstract

Advances in single cell whole genome sequencing enable profiling of the copy number state of thousands of cells with minimal sequencing bias across the genome. The Direct Library Preparation + technique is an whole genome amplification-free single cell whole genome sequencing method that achieves high throughput by fragmenting each cell’s genome and ligating sequencing adapters using a modified Tn5 transposase, and sequencing to less than 0.1x coverage. Despite recent advances in experimental approaches, data analysis of single cell whole genome sequencing lags behind and the existing methods are not optimized for the analysis of frozen samples with variable DNA preservation. Furthermore, existing tools predominantly rely on read depth ratio in predefined genomic bins to call copy number, making whole genome duplication unidentifiable. To address this, we introduce Songbird, a single cell whole genome sequencing copy number caller that is whole genome duplication sensitive, and outperforms existing tools both in breakpoints identification and true copy number detection. We demonstrate that Songbird is robust down to extremely low coverage, adaptable to a variety of genome versions (hg19, hg38, hs.1), and is extensible to other single cell whole genome sequencing methods that rely on Tn5 tagmentation to fragment the genome.

## 1 Introduction

Copy number variations (CNVs) are genome-wide changes observed across the majority of solid tumor types and often contribute significantly to the fitness advantage exhibited by primary tumor subclones [1]. CNVs can arise in many ways – for example, oncogene-driven failures in cytokinesis can result in whole genome doubling (WGD), enabling cells to lose significant genetic material without suffering haploinsufficiency of essential genes [2]. WGD is often one of the earliest changes in natural histories of gastric and esophageal adenocarcinomas [3]. Alternatively, failures in DNA damage response enzymes, or micronuclei formation cause rearrangements and genomic lesions which give rise to CNVs [4, 5]. Regardless of the mechanism behind their creation, CNVs may delete tumor suppressor genes, or produce multiple copies of oncogenic ones, potentially increasing the tumor fitness [6, 7]. The gain in fitness follows tumor evolution models, and recent work has linked copy number changes to immune evasion in ovarian cancer [8, 9]. While traditional in-line copy number changes are an intuitive driver of cancer evolution, the same processes driving copy number changes also creates structural variants (SVs) which contribute to tumor heterogeneity and plasticity [10].

Two such structural variants, breakage fusion bridges (BFBs) and extrachromosomal circular DNA (ecDNA), exhibit extremely high copy counts in cells with highly variable copies from generation to generation [11, 12]. These SVs, through their high count and highly variable composition, drive intratumoral heterogeneity and allow tumors to adapt rapidly to overcome treatments and immune surveillance [10, 13].

In both cases, the adaptability conferred to tumors by these SVs make them an important topic of research. Since these SVs can have wildly varying copy numbers, even within cells of the same phylogeny, it is important to study these structural variants on a single cell level. The tools best suited to studying this, single cell whole genome sequencing (scWGS), can be distinguished into three categories: PCR-, isothermal amplification-, and transposase-based [14]. PCR and isothermal amplification-based scWGS both rely on degenerate oligonucleotide primers (DOP) to randomly prime and amplify the genome [15, 16, 17]. Due to the reliance on primer binding and uniform amplification of DNA, the first two methods suffer from allelic dropout or uneven coverage of the genome, requiring high sequencing depth to reconstruct the copy number changes [18, 19].

Direct Library Preparation Plus (DLP+) is a Tn5 tranposase-based scWGS method that uses a robotic dispensing system to spot and lyse cells in nanoliter-scale wells of a 5184-well chip. The genomic DNA is converted into sequencing libraries in a sequential procedure that starts with simultaneous genomic fragmentation and library adaptor ligation facilitated by the Tn5 transposase. Following tagmentation, the Tn5 enzyme is deactivated and per-cell barcodes added to the genomic fragments via PCR amplification. Unlike DOP- or isothermal amplification-based methods, the Tn5 activity is randomly distributed across the genome, enabling accurate copy number recovery with shallow depth [20].

Furthermore, since library adapters are directly ligated to the genomic DNA, unique reads represent the original genomic fragments. However, due to the random adapter pairings on fragments, DLP+ has a maximum possible coverage of 0.5 times the cell’s overall ploidy [14]. Currently only a small family of tools dedicated to calling CNA from scWGS data exists.

The most straightforward of these, HMMCopy and SCOPE, rely on the read depth ratio of the binned read counts to fit copy number states. HMMCopy relies on a hidden Markov model to identify the transition point between copy number states [21]. Since the transition matrix in a hidden Markov model is finite, HMMCopy is unable to call copy numbers accurately in regions with high copy number state. SCOPE on the other hand fits a hierarchical model to the data, adjusting its copy number fitting for per-cell variations [22]. Since these tools rely only on binned read counts, they are unable to detect WGD events, as the ratio in read counts across bins follow similar distributions regardless of whether WGD has occurred.

Other tools attempt to address the WGD non-identifiability problem in a variety of ways. CHISEL and Alleloscope use single nucleotide polymorphism (SNP)-based phasing to estimate the cell’s average ploidy [23, 24]. However, since both tools rely on phasing, they need significant depth to accurately estimate copy number, and they would not detect perfect WGD where the ratio between alleles at heterozygous SNPs remain one-to-one. scAbsolute has attempted to address WGD detection in DLP+ data specifically by comparing overall read depth to local read density. It also relies on a hidden Markov model to identify copy number breakpoints, but uses an expectation maximization algorithm to fit the final copy number [25]. Interestingly, even though the unique reads in DLP+ are derived from unique fragments of the genome, no tool has been able to use that information to estimate true ploidy, largely due to the low coverage of the DLP+ data itself. Furthermore, the Tn5 enzyme leaves nine nucleotide overlaps in adjacent reads, providing extra information about adjacent fragments from the same allele.

Leveraging this unused information, we present Songbird, a Haar wavelet based, absolute ploidy sensitive copy number caller adaptable to any Tn5-based whole genome sequencing method (Figure 1). In contrast to other tools, Songbird uses unbalanced Haar wavelets to sensitively identify the breakpoints in the genome [26]. In addition, the tool leverages the unique features of Tn5-based scWGS to both estimate the average ploidy of the cell and the amount of the genome available for sequencing initially. Lastly Songbird uses a likelihood ratio maximization algorithm to estimate the true copy number given the segmented regions and estimate of true ploidy. We benchmark Songbird against simulated scWGS data, existing DLP+ datasets and ploidy ladders, and show that Songbird more accurately captures high copy number events and estimates ploidy better than competing tools. In addition, Songbird is extensible to data aligned to hg19, hg38, and hs.1 versions of human genome.

**Figure 1:**
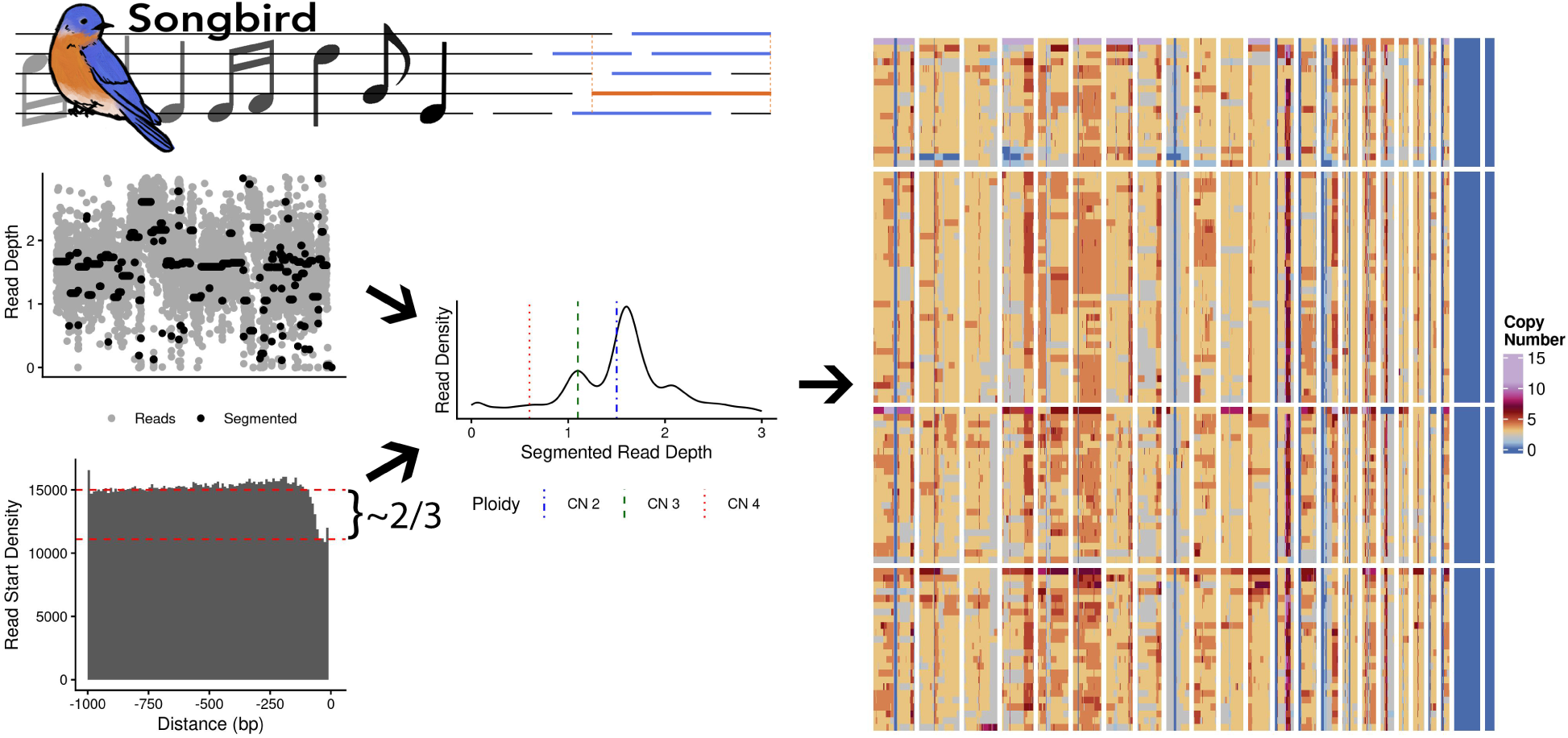
For each cell sequenced by DLP+, Songbird simultaneously estimates the copy number breakpoints, and the cell’s overall ploidy. These two pieces of information are integrated together to fit copy number states which maximize fit, and refines the copy number calls by sharing estimated ploidy information across all cells. The example cells are triploid, so the overlapping read ratio for ploidy estimation approximates 2/3.

## 2 MATERIALS AND METHODS

### 2.1 Patient Information

Endoscopic and surgical samples were collected at the Cambridge University Hospitals NHS Trust (Addenbrooke’s Hospital) from esophageal adenocarcinoma patients undergoing cancer resection. The study was approved by the Institutional Ethics Committees (REC 07/H0305/52 and 10/H0305/1), and all subjects gave individual informed consent.

### 2.2 Single Cell DNA Sequencing

#### 2.2.1 Tissue Culture

The immortalized hTert cell lines were derived from a male lymphoblastoid cell line. The hTert and HEK293T cell lines were cultured at 37*^◦^*C in 5% CO2, and grown in RPMI-1640 or DMEM respectively. Both media are supplemented with 2mM L-glutamate and 10%FBS. Cells were passaged every 2-3 days after achieving approximately 90% confluency. To passage, the cells were washed with 1x PBS, detached using 0.05% Tripsin and reseeded at 1:5 split, and cell passage number was recorded.

#### 2.2.2 DLP+ Sequencing

DLP+ was performed in keeping with previously published protocols [20]. In order to create the sequencing ladder, a mixture of cells were placed into each well such that the total genomic content ranges from 2N to 10N. For each of the cell mixtures (1x,2x,3x,4x hTERT, 1x HEK293T, 1x hTERT and 1x HEK293T, 2x hTERT and 1x HEK293T, 1x hTERT and 2x HEK293T, 2x hTERT and 2x HEK293T), we spotted 100 wells. Prior to cell spotting, Illumina-compatible primers containing unique cell barcodes are spotted into 72×72 array. After cells spotting each well is loaded with QIAGEN protease, Viagen DirectPCR cell lysis buffer, and the cells heat lysed at 50*^◦^*C for 60 minutes followed by buffer inactivation at 75*^◦^*C, 15 min. The tagmentation buffer is then added and incubated at 55*^◦^*C for 10 minutes. The Tn5 reaction was stopped by neutralization via Qiagen Protease at 50*^◦^*C for 10 minutes followed by an additional heat inactivation of the protease (70*^◦^*C, 15 min). After deactivation of the Tn5 enzyme, the samples underwent 11 cycles of PCR and the resulting amplified DNA were pooled from the individual wells and sequenced on an Illumina NovaSeq 6000 S4 flow cell to produce 150bp paired end reads.

### 2.3 Data Preprocessing

The ploidy ladder and the esophageal adenocarcinoma sample were processed using the *mondrian* pipeline (https://github.com/mondrian-scwgs/mondrian); reads were trimmed using *TrimGalore v0.6.6*, and quality checked with *FASTQC v0.11.9.* Trimmed reads were then assigned to a read group based on the trimmed adapter sequence and aligned to the GRCh38 genome with decoy and alternative contigs and screened for contamination using *FastQScreen v0.14.0*. The per cell bam files were filtered to retain high quality (phred score greater than 30), primary alignments mapped to the 22 autosomes and two sex chromosomes. The filtered bam is deduplicated using *samtools (v1.16) fixmate* and *markdup* functions before using *bedtools (v2.30.0) bamtobed* to generate the bedpe file that identifies genomic regions spanned by each fragment.

To prepare the SNV data for *Alleloscope*, we followed the tutorial provided on their github repository. This involved producing phased SNV calls from the bulk bam, converting the calls to a cell-by-position matrix using *Vartrix* (https://github.com/10XGenomics/vartrix), and generating a counts matrix in the same format. *Alleloscope*, *SCOPE*, and *scAbsolute* were run in keeping with their respective documentation. *HMMCopy* was run in a small optimization loop based off the *mondrian* pipeline. This involves multiplying the GC and mappability corrected read density against a series of multipliers which will convert the read density into integer states. The multiplier value which minimizes the number of half copy calls (i.e., copy numbers 1.5, 2.5, 3.5 etc.) is retained as the true multiplier.

### 2.4 Simulated Read Generation

We use *CINner* [27], an algorithm for modeling the evolution of chromosomal instability, to create simulated copy number profiles. Each simulation starts with 1,000 diploid cells, and the cell division probabilities are calibrated such that the total population size remains constant on average. Each cell division contains a missegregation of a randomly selected chromosome with probability *p*_misseg_ ∈ {0.02, 0.05, 0.08, 0.2}. The copy number profiles observed after an average of 300 generations are then recorded.

We then simulate cell- and bin-specific readcounts. For a given cell, let {*n_j_, g_j_*} be the simulated copy number and GC content of bin *j*. We assume a total of *r* = 4 × 10^6^ reads per cell, following Gamma distribution per bin with scale *σ* = 0.02 (approximated from previous DLP+ data). We simulate read count *r_j_* for bin *j* as:

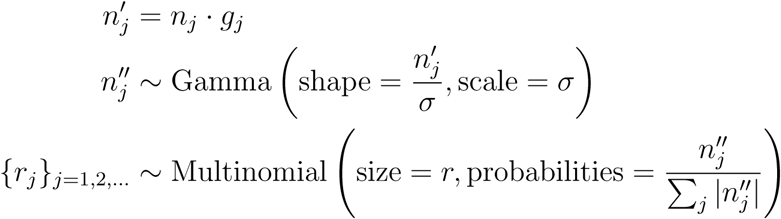

The inferred copy numbers based on {*r_j_*} are then compared against ground-truth copy numbers {*n_j_*} to determine accuracy.

### 2.5 Songbird

Songbird is composed of four parts – breakpoint identification, ploidy estimation, subclone/whole genome duplication detection, and finally integer fitting.

#### 2.5.1 Copy number breakpoint identification

We use *QDNASeq* to load in reads from the bam file, bin them, and perform read depth correction per cell [28]. We adjust the read depth with respect to mappability and GC content using correction factors calculated using only autosome aligned reads, but applied to the whole genome. In order to identify breakpoints where the true copy number state changes in the corrected read density, we use the unbalanced Haar wavelet transform [26]. This algorithm fits asymmetrical Haar wavelets to time series data. It does so by splitting the data at the most probable point where the average value after the breakpoint is distinct from the average value before the breakpoint and assigns a score to the split. The algorithm then recurses on the time span before and after the breakpoint separately, repeating until the data can not be split anymore, or all values are identical. Splits with support greater than a threshold are retained. In this case, we treat the entire genome as the time series data, ordered by chromosome and coordinate, and the threshold is set to the median absolute deviation of the read depth, as recommended by the paper.

#### 2.5.2 Ploidy and Available Genome Estimation

The genomic coordinates covered by each fragment are loaded in from the bedpe file and further filtered to remove artifactual and short fragments. Artifactual fragments are defined as fragments which share a start or end coordinate with another longer fragment and oriented in the same direction (e.g. for both fragments, the forward sequencing read aligned to the Watson DNA strand and reverse read to the Crick DNA strand). We apply a user-defined minimum fragment length cutoff; typically 50 nucleotides for Illumina short read sequencing. We also remove the first 10 nucleotides from each fragment span to correct for the overlap generated by Tn5 tagmentation between adjacent fragments.

In order to estimate the ploidy *n* of given cells we work with pre-filtered sequencing fragments. As mentioned above, we know that, the DLP+ chemistry ensures that the spans covered by each unique fragment represent a unique allele. If we choose a sequenced fragment and designate it as the *reference fragment*, we know that there are *n* − 1 copies of the genome available to generate fragments at the genomic coordinates overlapping this reference fragment, and *n* copies of the genome wherever the reference fragment is not. Given a known minimum fragment length, we know that fragments starting less than that length upstream of the reference fragment must overlap the reference fragment, therefore are from unique alleles. We designate such fragments as *overlapping fragments*. In contrast, fragments starting significantly upstream (greater than the maximum length of all fragments) of our reference fragment are drawn from all *n* copies of the genome, as they should never overlap the reference fragment. We designate these as *upstream fragments*. In practice, we look for the upstream fragments in a variable window that starts at the maximum read length before the start of the given reference fragment and ends at the minimum read length before the start of the reference fragment. Since the longest fragments rarely exceed 1.5 kilobases in Illumina short read sequencing, the upstream and overlapping fragments are likely drawn from regions of the genome with similar accessibility.

With these assumptions for each reference fragment with the start coordinate *a*, we calculate the count of overlapping fragments in the window (*a* − min(fragment.length)*, a*) and the count of the upstream fragments in the window (*b, a* − max(fragment.length)), where *b* is a genomic location significantly upstream of *a*. To account for the different length of the overlapping and upstream windows, we normalize the counts by their respective window size. The overlapping fragment start density and upstream density is converted to the estimated ploidy using the equation:

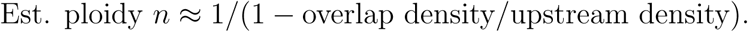

For extremely long fragments, such as those generated by PacBio sequencing, the kilobase fragment lengths mean that long fragments could stretch into regions of the genome which were not similarly preserved or deacetylated as the reference region. Therefore, we have to enforce an upper limit on the fragment lengths, which in our limited experience is approximately eight kilobases.

#### 2.5.3 Subclone Identification, Ploidy Correction, and Whole Genome Duplication detection

To help correct the noisy ploidy estimates we use information from all cells and identify subclones with similar copy number states and pool the ploidy estimates to get a more accurate value. Subclones are identified by clustering the positions of the copy number breakpoints between cells. In more detail, we find breakpoints that are observed in at least 10% of cells, apply a Gaussian smoothing kernel to minimize the difference between breakpoints that are offset by one bin, and construct a graph between the *k* nearest (Manhattan distance) neighbors to each cell. This graph is entered into the Phenograph algorithm to identify subclones [29]. Within each subclone, we gather estimated average ploidies (greater than 0 and less than 10) from cells with more than 50,000 reads and fit a *k* = 2 K-means clustering model on the values. If the clusters are sufficiently compact and the centroids are close to a 1:2 ratio, cells are assigned WGD status based on their cluster membership.

#### 2.5.4 Integer fitting

We use the average ploidy estimate to propose a targeted range of possible average integer ploidies for the cell. For each of the tested average ploidies we calculate a reads per copy number (RpCN) value using the median segmented value. Since the median segmented value only provides an approximation of the true RpCN, we need to optimize each proposed RpCN prior to evaluating the one with best fit (Figure 4B).

To score the RpCNs, we calculate their likelihood using a model where read density per bin is generated from a normal distribution centered about the true RpCN times the true copy number state with some noise. We can make a conservative estimate of the magnitude of the noise by looking at the read density error with respect to the segmented read states. Finding the RpCN by just maximizing the likelihood of this model alone would always favor a lower RpCN, as a smaller RpCN will maximize the distribution densities by proposing more overlapping distributions. To account for this we also generate a null read distribution for each cell drawn from a normal distribution centered at the overall read density average using the same noise calculated above. This likelihood ratio between the RpCN applied to the segmented data and the null data is then used to both optimize each proposed RpCN, and the RpCN with the largest likelihood ratio is selected.

## 3 RESULTS

### 3.1 Tn5-based scDNA libraries contain artifactual reads that bias the frequency of read overlaps

Since DLP+ generates fragments derived from unique copies of the genome, simply observing the patterns of overlapping fragments would be an intuitive way to call the copy number profile. Ostensibly the maximum fragment depth at any location would indicate the true copy number state of that region of the genome. We observed two difficulties with this approach. First, the shallow nature of DLP+ sequencing meant the maximum fragment depth significantly underestimated the true copy number of high copy number regions. Second, even after excluding problematic genomic regions such as those found on the ENCODE exclusion list [30] and only retaining regions with high mappability scores, we observed a significant number of regions in the genome where the overlap state incremented in steps of two or higher indicating multiple reads starting or ending at the same coordinates. When compared to a random simulation read generation, we observe a significantly higher proportion of two state transitions than expected (Figure 2A). Since we removed classically defined (the same start and end coordinates) duplicate fragments prior to our bedpe file generation, this indicated that there were a large number of fragments which started or ended at the same position but were of different lengths. Furthermore, our data was generated with a diploid, hTERT overexpressing cell line with no copy number variation outside of chromosome 20 (Supplementary Figure 1). Therefore the maximum expected depth should not exceed 2, which is clearly violated by these fragment pairs. Interestingly, a large majority of these pairs had identical genomic orientations, outnumbering pairs with opposite orientations in a nearly 10:1 ratio (Figure 2B).

**Figure 2:**
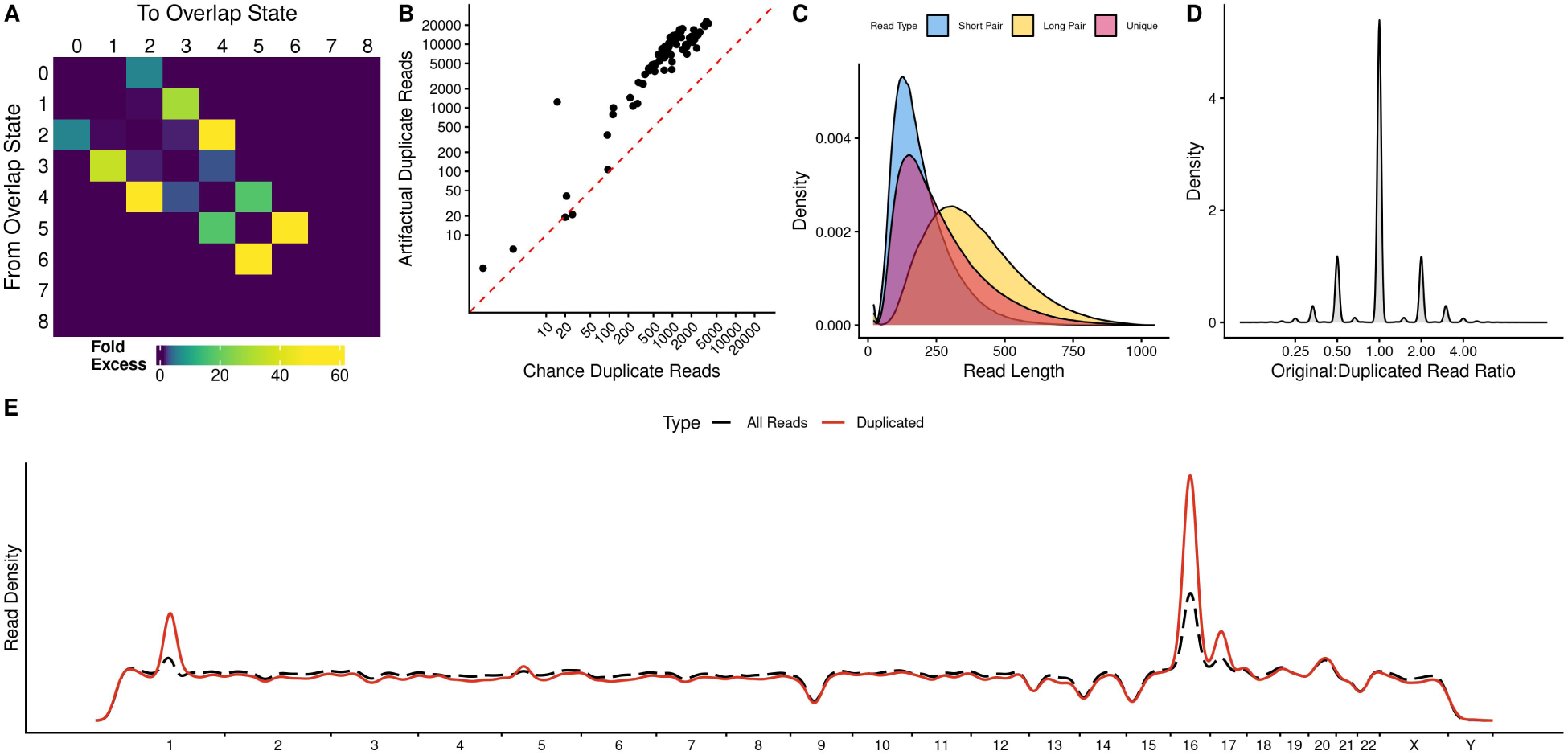
(A) Heatmap of the fold excess in observed transitions in 100 hTERT Cells vs a simulation. A high fold excess proportion of 0*>*2 transitions indicates that there are more reads starting at the same coordinate than expected by random read placement (B) Scatterplot comparing the number of artifactual duplicate fragments (same start or end and same orientation) with the number of chance fragments (same start or end, but different orientations) in 100 diploid hTERT cells. (C) Fragment length distributions for the shorter (blue) and longer (yellow) member of the artifactual fragment pairs compared to the unique fragment length distribution (red). (D) Ratio of longer (original) to shorter (duplicated) fragment counts in the original bam prior to standard duplicate (same start, end and orientation) read removal. (E) Density across the genome for artifactual and unique fragments calculated over 100 diploid hTERT cells.

Looking at the fragment pairs themselves, we observe that the shorter fragment in these potentially artifactual pairs were shorter than the unique fragment length distribution, while the longer fragment were longer (Figure 2C). Further investigation of these pairs prior to duplicate fragment removal showed that these fragments pairs are overwhelmingly amplified in a 1:1 or 1:2 ratio, hinting that the process which produces these fragments occurs within or prior to the first cycle of PCR amplification (Figure 2D). While there are some regions where these fragments are more common than others, the density of these potentially artifactual sequenced fragments largely mirrors the distribution of all fragments generated in these cells (Figure 2E). Furthermore, any deviations between all fragment and artifactual fragment density seems to be a function of genome version as they shift in location depending on the genome that the data is aligned to (Supplementary Figure 2). Since they are a minority of the total fragment count (*<*1%) but constitute a significant proportion of overlapping fragments, we discard the shorter fragment from each artifactual pair. Discarding these reads improves ploidy estimation performance.

### 3.2 Songbird accurately recalls copy number breakpoints in simulated and real scDNA data

In order to validate the breakpoint sensitivity of the unbalanced Haar wavelet algorithm we simulated reads from 4000 cells using CINner [27] with increasing rates of chromosomal instability (Figure 3A). To test the tool, we fit Songbird and HMMCopy to each cell and compared the break points and overall fitting to the ground truth copy number state. Breakpoint recovery was calculated by calculating the Jaccard index between the coordinates of the estimated and true breakpoints. Since even a one-bin offset will lower the Jaccard index, this yields relatively low scores, but Songbird still recovered a much higher proportion of true breakpoints compared to HMMCopy (Figure 3B). Both Songbird and HMMCopy provide a segmented read depth value prior to final copy number fitting. Scaling these read depth values to the true copy using each simulated cell’s actual read per copy number ratio demonstrates that Songbird’s segmented states mirror the true copy number more closely than HMMCopy (Figure 3C). In order to test Songbird’s breakpoint estimation in real data, we generated a ploidy ladder by spotting a DLP+ nanowell array with combinations of hTERT and HEK cells such that the expected ploidy per well ranges from 2 to 6. As we downsample this ploidy ladder, the unbalanced Haar segmentation is consistent, maintaining high correlation with the initial segmentation down to 0.005x coverage, or approximately 100k reads per cell if using an Illumina short read sequencer (Figures 3D, E).

**Figure 3:**
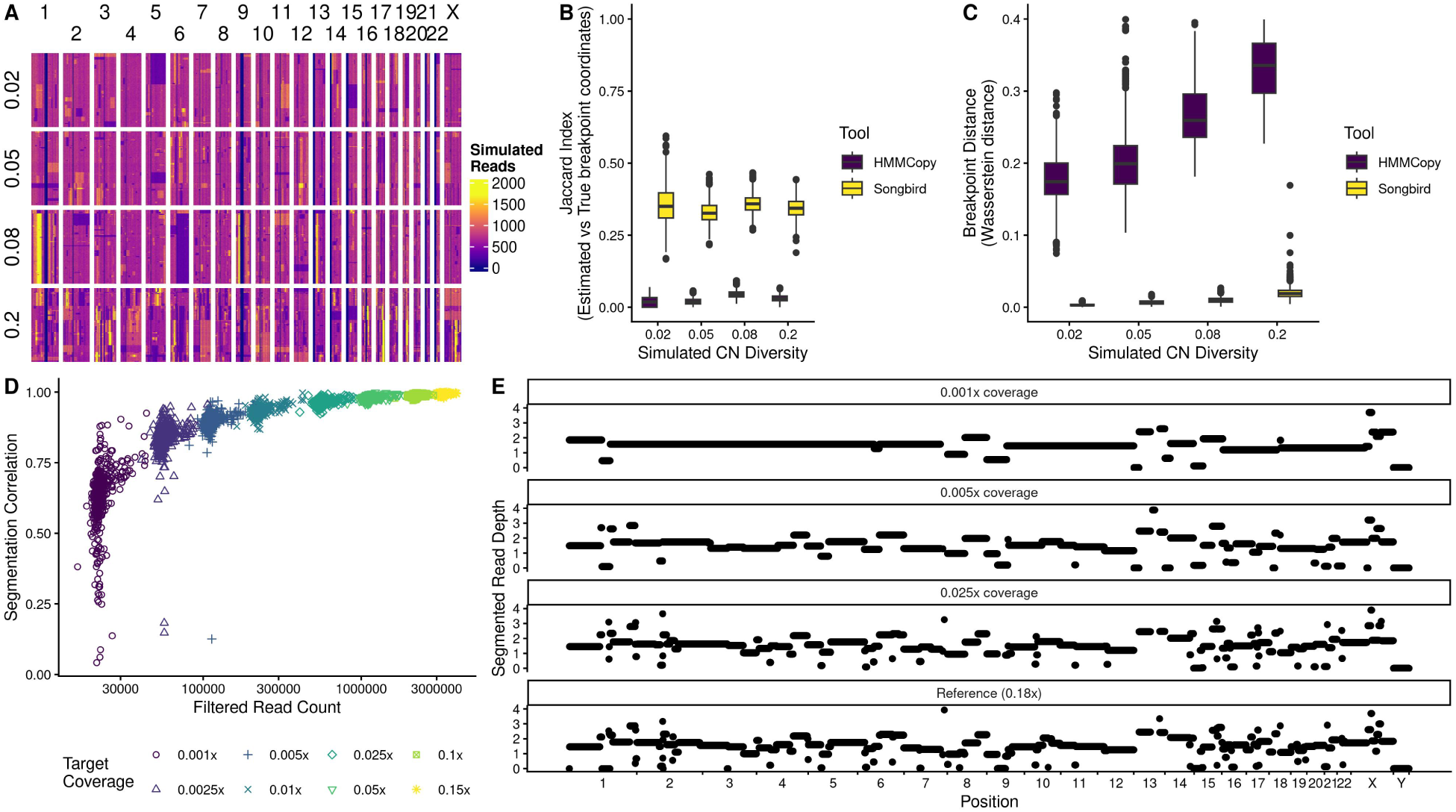
(A) Heatmap of simulated read depth in reads generated by CINner with increasing missegregation rate parameters (B) Jaccard Index scoring the recovery of the simulated breakpoints in the simulated data. (C) Wasserstein Distance between the segmented state and true copy number in the simulated data. The segmented states were scaled by each cell’s true reads per copy number ratio prior to distance calculation. (D) Correlation with initial segmented states across the sequenced ladder across downsampled cells. Cells with lower coverage than the target coverage were omitted. (E) Qualitative comparison of the segmented state across downsamplings for 1 HEK293T Cell.

### 3.3 Songbird accurately estimates total cell ploidy

In order to estimate cell ploidy, we take advantage of the fact that unique fragments generated in a DLP+ run are all derived from exclusive fragments of the original genome. Given this information, if we observe a reference read in a region with *n* copies of the genome, we know that there are only *n*−1 copies left to draw overlapping reads from. In practice this calculation is not straightforward, since a long reference fragment can be overlapped by multiple shorter fragments derived from the other alleles. To resolve this problem we enforce a minimum fragment size cutoff *f*_short_. This enables us to look at a window of *f*_short_ nucleotides upstream of our reference fragment and know that the observed fragments starting in that window must come from different alleles and must overlap our reference fragment. If we look more than the longest read length, *f*_long_, upstream of our reference fragment, we know that the reads starting in that window do not overlap the reference fragment and are derived from *n* copies of the genome. The density of fragments overlapping our reference fragment divided by the density of fragments far upstream of our reference fragment should approach (*n* − 1)*/n* with sufficient coverage. Following simple algebra, the ploidy *n* of a cell can then be estimated as *n* ≈ 1*/*(1 − overlap density*/*upstream density).

Songbird provides a practical algorithm (see Methods) for the estimation of a cell’s average ploidy by treating each read as the reference fragment, repeating this calculation, and averaging the ratios together. The initial removal of artifactual reads is important for the accuracy of this method, which we validated by spotting multiples of diploid hTERT and triploid HEK293T cells into individual wells of DLP+ chips. We were able to estimate true copy number of the spotted ladder, which ranged in copy number from 2 to 10, reliably up to copy number 6 (Figure 4A). The accuracy of caller was independent of the version of human genome assembly (Supplementary Figure 3A). Crucially, since the reference hTERT cell line is free of copy number alterations, our estimator works in the absence of breakpoints and can clearly distinguish diploid and tetraploid-like (two cells) without relying on variations in copy number to distinguish ploidy. Furthermore, Songbird’s ploidy estimator produces consistent ploidy estimates using smaller bins sizes (down to 50 kbp, Supplementary Figure 3B) and in high CNA (average ploidy 5) cells down to 0.01x coverage, highlighting its robustness (Supplementary Figure 3C).

**Figure 4:**
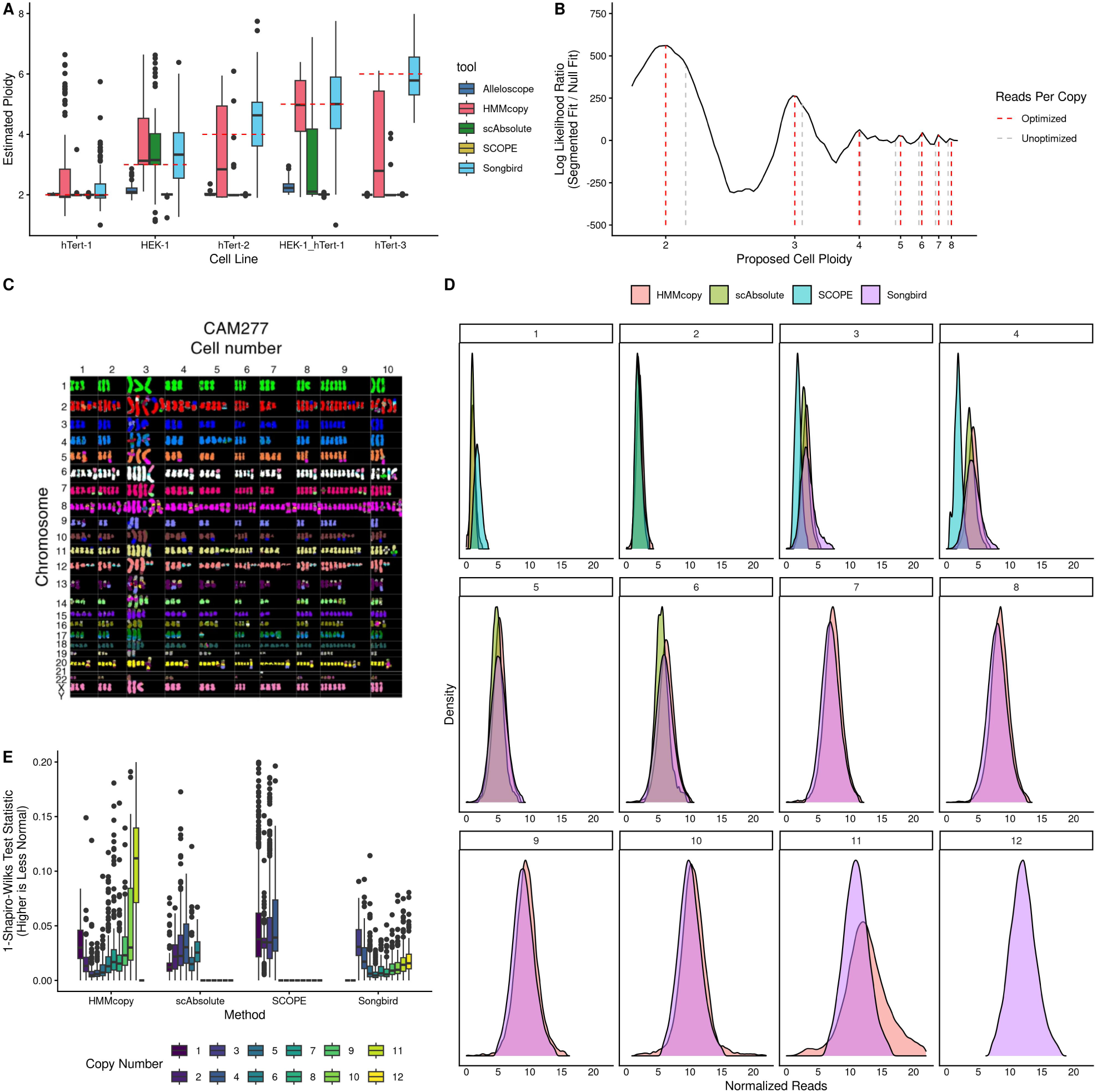
(A) Raw ploidy estimates across our cell ploidy ladder. (B) Log Likelihood Ratio fit for a range of reads per copy number (RpCN) ratios in a HEK293T cell. Grey lines indicate the initial RpCN ratio proposed using the median segmented state and red lines indicate the optimized ratio. (C) Karyotyping of 10 CAM277 organoid cells. Figure used with permission from Li et al 2018 [31]. (D) Qualitative comparison of the distribution of reads assigned to each copy number across 581 cells sequenced from the CAM277 organoid. (E) Quantative test of normality for the reads assigned to each copy number. The test statistic was subtracted from 1 so that more normal distributions have a lower score.

Despite its simplicity and accuracy in high quality data, the stochastic nature of scDNA sequencing introduces uncertainty in ploidy estimation, as evidenced by the large area covered by the boxplots in Figure 4A. In order to gain additional knowledge and improve accuracy of ploidy calling, we identify clusters of cells with similar breakpoint patterns. To do so, we apply phenograph clustering on the common breakpoints and average the clustered cell ploidies to estimate corrected ploidy in each cell and flag WGD cells.

In order to call copy number, we aim to find the correct reads per copy number ratio (RpCN) for each cell. However, since our ploidy estimate is noisy, we test a set of RpCN values that correspond to true copy number states around the ploidy estimate. These are derived by dividing the median segmented read depth by each proposed average ploidy. Since the median read depth may not be a perfect multiple of the true RpCN for that cell, we optimize these RpCNs for each proposed ploidy for the maximum likelihood ratio compared to a null distribution generated from each cell (Figure 4B). To validate Songbird’s copy number fitting, we used DLP+ sequencing data from a previously published esophageal adenocarcinoma organoid as it had both a significantly varied copy number profile, and karotyping to validate the true copy number states (Figure 4C) [31]. Compared to SCOPE, scAbsolute, and HMMCopy, the reads assigned to each copy number by Songbird were more normally distributed, indicating better fit (Figure 4D). This is quantified in Figure 4E by subtracting Shapiro-Wilks test statistic from 1 so that 0 is perfectly normally distributed. Copy numbers that were not called by the various methods were given a zero score. Lastly, our ploidy estimator is extendable to other Tn5 based scWGS methods such as Lianti [32], and SmoothSeq [33] (Supplementary Figure 4A). Furthermore, it can accurately assign copy number states to cell cycle sorted cells (Supplementary Figure 4B) [20] and the analyzed previously published DLP+ data demonstrate its universality (Supplementary Figure 4C and 4D) [25].

### 3.4 Recapitulating single cell ecDNA Dynamics with Songbird

While most scWGS copy number callers segment the genome with 500kbp bins, Songbird is able to segment the genome down to 30kbp bins with relatively high fidelity with respect to the 500kbp segmentation (Supplementary Figure 3B). While the arbitrary bin starting and end points do not recapitulate the actual genomic breakpoints of structural variants, they can approximate them (Figures 5B, C). Previously, we collected Oxford Nanopore long read sequencing from the same esophageal adenocarcinoma organoid and we used it to reconstruct the structure of an ecDNA and a BFB in that sample [34]. Songbird run at 50 kbp bin size effectively approximated the ecDNA breakpoints, and the bins identified displayed the expected ecDNA dynamics across cells (Figure 5D). Namely, while the copy number of the ecDNA in each cell varies significantly, the copy number ratio between fragments within the ecDNA are constant across cells, regardless of the total ecDNA copy number state. In contrast, the BFB does not show consistent ratios across overlapping bins, consistent with the underlying processes driving inheritance of these structural variants (Figure 5E)

**Figure 5:**
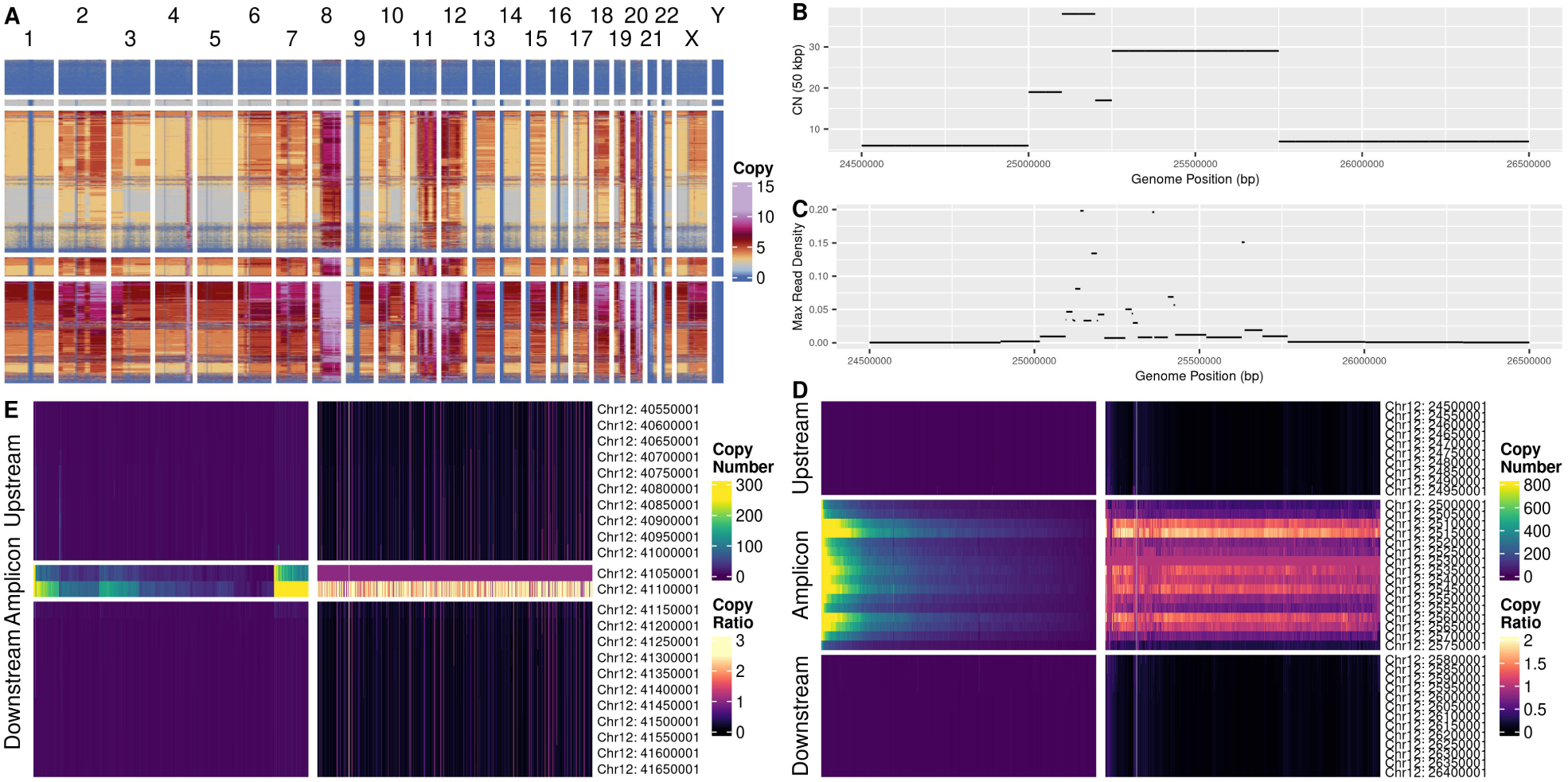
(A) Full Copy Number profile of the CAM277 organoid broken into subclones. (B) Copy number called on 50kbp bins in a region with a known ecDNA in the organoid. (C) Maximum read depth across the known ecDNA fragment intervals. Intervals for ecDNA reconstruction were generated using Oxford Nanopore long read sequencing [34]. (D) Copy number profile and bin ratios for the 50kbp bins overlapping the ecDNA interval. (E) Copy number profile and bin ratios for 50kbp bins overlapping a separate, known BFB interval. The BFB interval was reconstructed similarly to the ecDNA intervals [34].

## 4 DISCUSSION

Here we present Songbird, a wavelet-based, WGD-aware copy number caller for scWGS methods which rely on tagmentation for the initial cleavage and library adapter deposition. We demonstrate that Songbird, using the unbalanced Haar wavelet transform, is able to identify accurately copy number breakpoints in simulated DLP+ data, and reliably recover those breakpoints in cells with as little as 0.005x coverage. Furthermore, Songbird’s novel ploidy calling algorithm is able to recover the true ploidy of cells up to an average ploidy of 6, and consistently estimate the correct ploidy in cells with anywhere from 0.0025x to 0.01x coverage depending on the cell’s average ploidy (Supplementary Figure 4C). In addition, we demonstrate the ploidy calling algorithm’s versatility to a wide variety of protocols, successfully recapitulating the true ploidy from several other Tn5 based scWGS sequencing methods. Lastly we demonstrate the accuracy of Songbird’s final copy number fitting algorithm, and demonstrate its ability to recapitulate the expected single cell copy number dynamics of an already known ecDNA.

In developing Songbird, we have identified and partially characterized artifactual fragment pairs that seem to have interesting properties: these fragment pairs occur far more often than expected by chance, violate the known maximum ploidy in cells, and are largely duplicated in a 1:1 or 1:2 ratio. The first two observations hint at their artifactual nature, and the last seems to indicate that these fragments are generated prior to PCR amplification or within the first cycle. The only steps prior to PCR amplification in the DLP+ protocol are adapter tagmentation, and Tn5 neutralization. As a result, we suspect that these reads are generated from an incomplete Tn5 library adapter ligation, where one pair of the Tn5 dimer successfully cleaves one strand of an already fragmented piece of genomic DNA, while the other one fails to do so. This successful, failed cleavage pairing would produce two single stranded DNA fragments, one longer one representing the original, complete Tn5 transposase action, and a shorter one from this partially complete Tn5 cleavage. Since the adapter loading on the Tn5 dimer is random, we expect that we only observe half of these events, with the other half depositing same orientation primers and failing to amplify the short fragment.

We have not performed a biochemical exploration of these events due to their relatively rare occurrence despite their outsize impact on ploidy estimation. These events could be caused by anything from Tn5 cleavage events interrupted by the neutralization buffer, to incomplete neutralization, to even defective Tn5 monomers which are able to successfully dimerize with a functional monomer.

While Songbird seems to perform well, further work can be done to improve the noisy ploidy estimator. Currently, Songbird uses the average overlapping and upstream read densities to estimate true ploidy, but a more sophisticated Poisson or binomial sampling model may be able to more accurately capture the true copy number state. Furthermore, Songbird currently treats cells as independent events with very little shared information from cell to cell beyond clustering and ploidy estimate correction. A more holistic model that uses information across multiple cells would be useful. This would, for example, dramatically improve the accuracy of copy number calls at focal amplifications. Currently, at coarse bin sizes, the copy numbers from focal amplification events are estimated with only a single bin of information, which adds noise into the true copy number estimation since those reads are subject to the same sampling error which makes segmentation a necessity.

## 5 CONCLUSION

Single Cell Whole Genome Sequencing is essential for understanding tumor heterogeneity. The highly varied copy number landscape of cancer cells dramatically improves the fitness of these tumors, allowing them to escape local immune surveillance, evade cell intrinsic antitumor processes, and even gain resistance to therapeutics. The copy number landscape of these tumors is still to be studied, but in many cases seems to be enabled by whole genome doubling, making detection of these events critical. Furthermore, key structural variants such as breakage fusion bridges and ecDNA have been shown more recently to be critical mechanisms allowing tumors to resist treatment. While critical, these structural variants tend to be composed of small fragments, far smaller than the standard 500kbp bin size typically used in existing scWGS copy number calling tools.

Songbird is uniquely positioned to help explore the copy number dynamics of tumors at a single cell resolution in a manner which can detect these critical biological processes. Songbird has mostly been tested on DLP+ scWGS data, but has shown promising performance on other Tn5 based scWGS methods and even with PacBio long read sequencing. This indicates that Songbird may be adaptable to future Tn5 based scWGS methods.

## 6 DATA AVAILABILITY

Songbird is available as an R Package at https://github.com/GastroEsoLab/Songbird. Sequencing files for the hTERT/HEK ladder are available upon request. Files for the scAbsolute Ladder were retrieved from the European Nucleotide Archive project PRJEB61928. Sequencing files for Lianti and SmoothSeq were retrieved from NCBI bioproject PRJNA379710 and PRJNA633502 respectively.

Sequencing files for the esophageal adenocarcinoma organoid are available under restricted access.

## ACKNOWLEDGEMENTS

We thank Jennifer Jones for designing the banner in the Graphical Abstract.

This work was made possible by the MacMillan Family Foundation as part of the MacMillan Center for the Study of the Non-Coding Cancer Genome at the New York Genome Center.

---

**Supplementary Figure 1:**
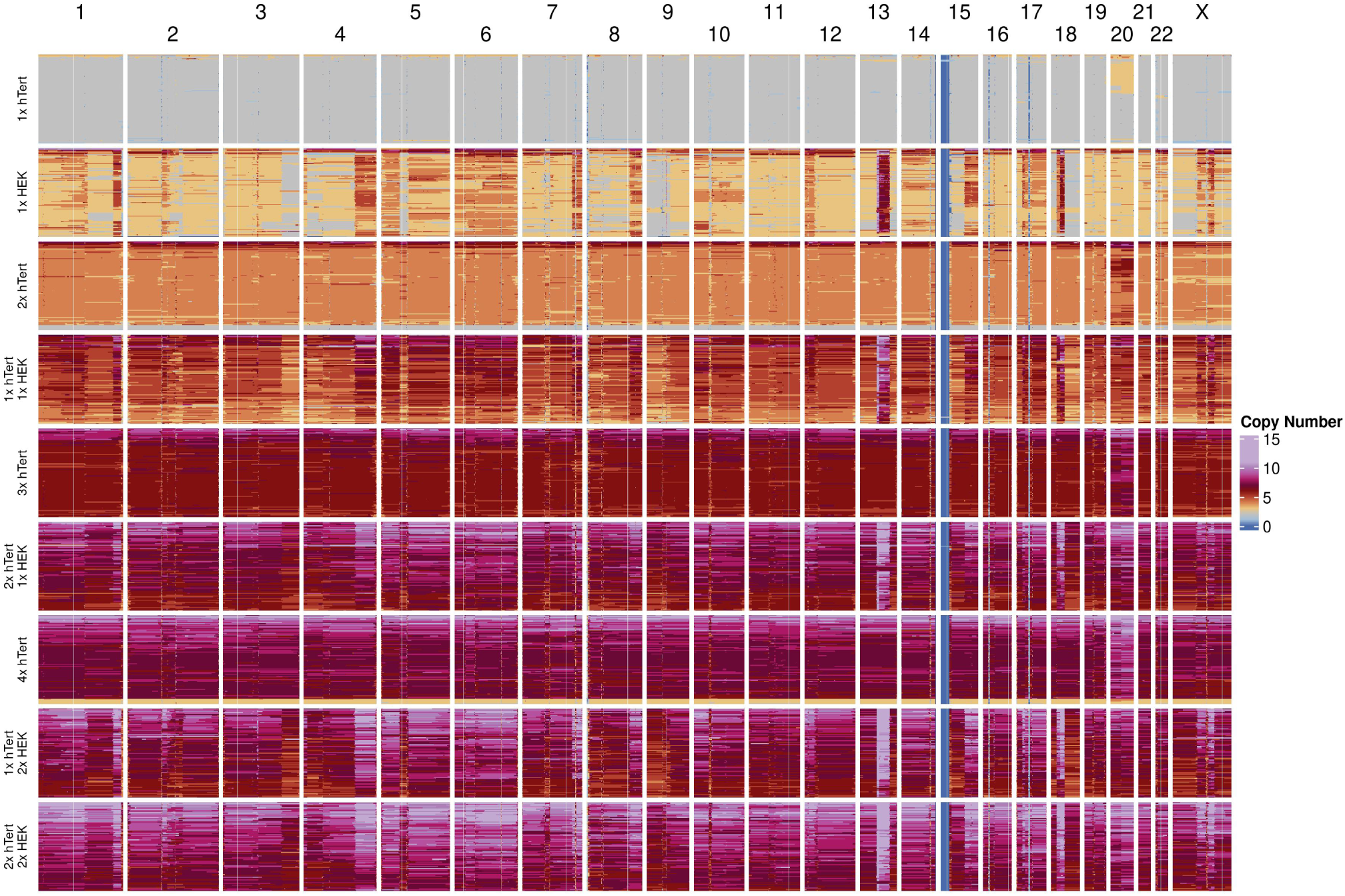
Assigned Copy Number profile for the ploidy ladder.

**Supplementary Figure 2:**
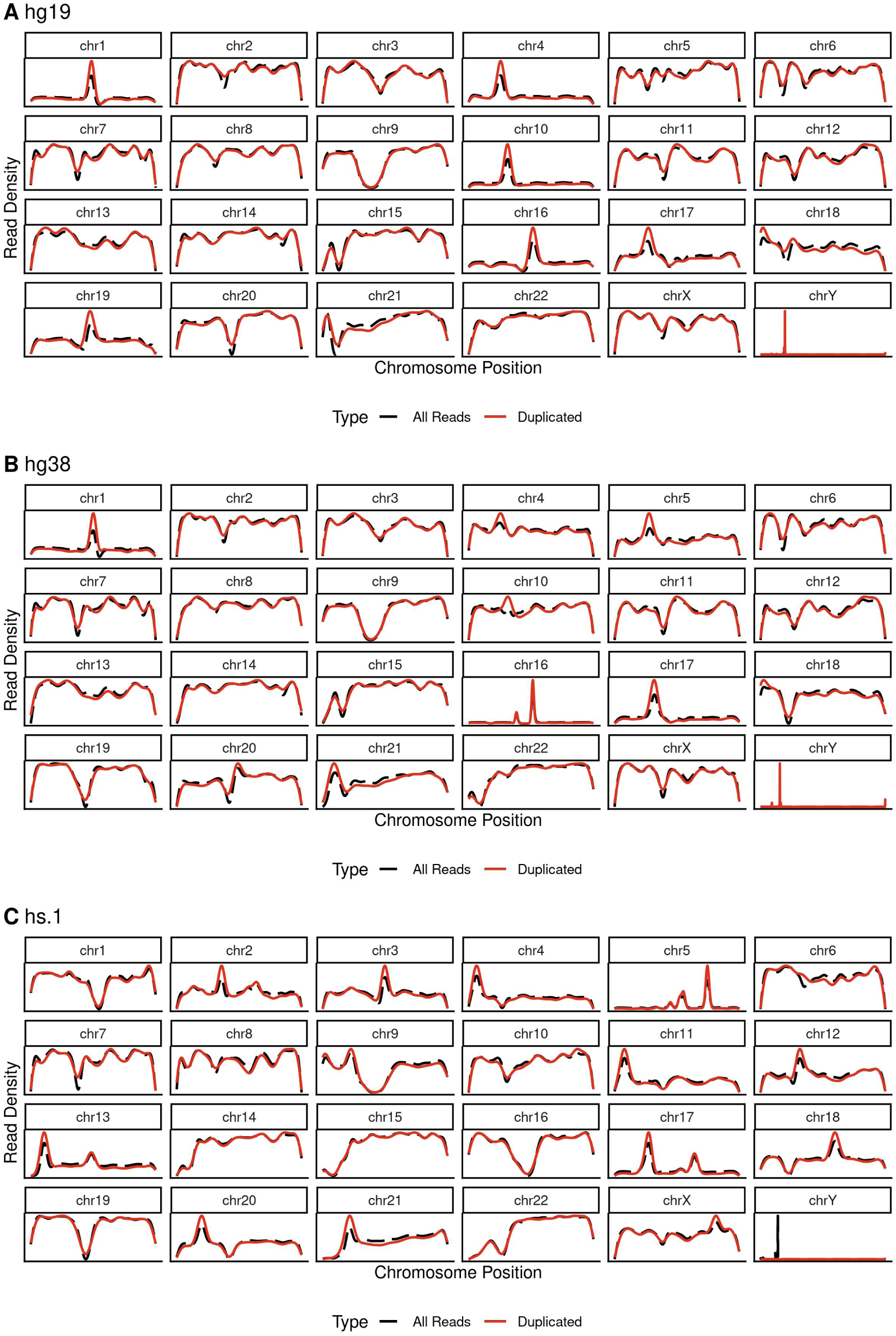
Density of artifactual reads (red) vs all reads (black dashed) for 100 diploid cells aligned to (A) hg19, (B) hg38, (C) hs.1.

**Supplementary Figure 3:**
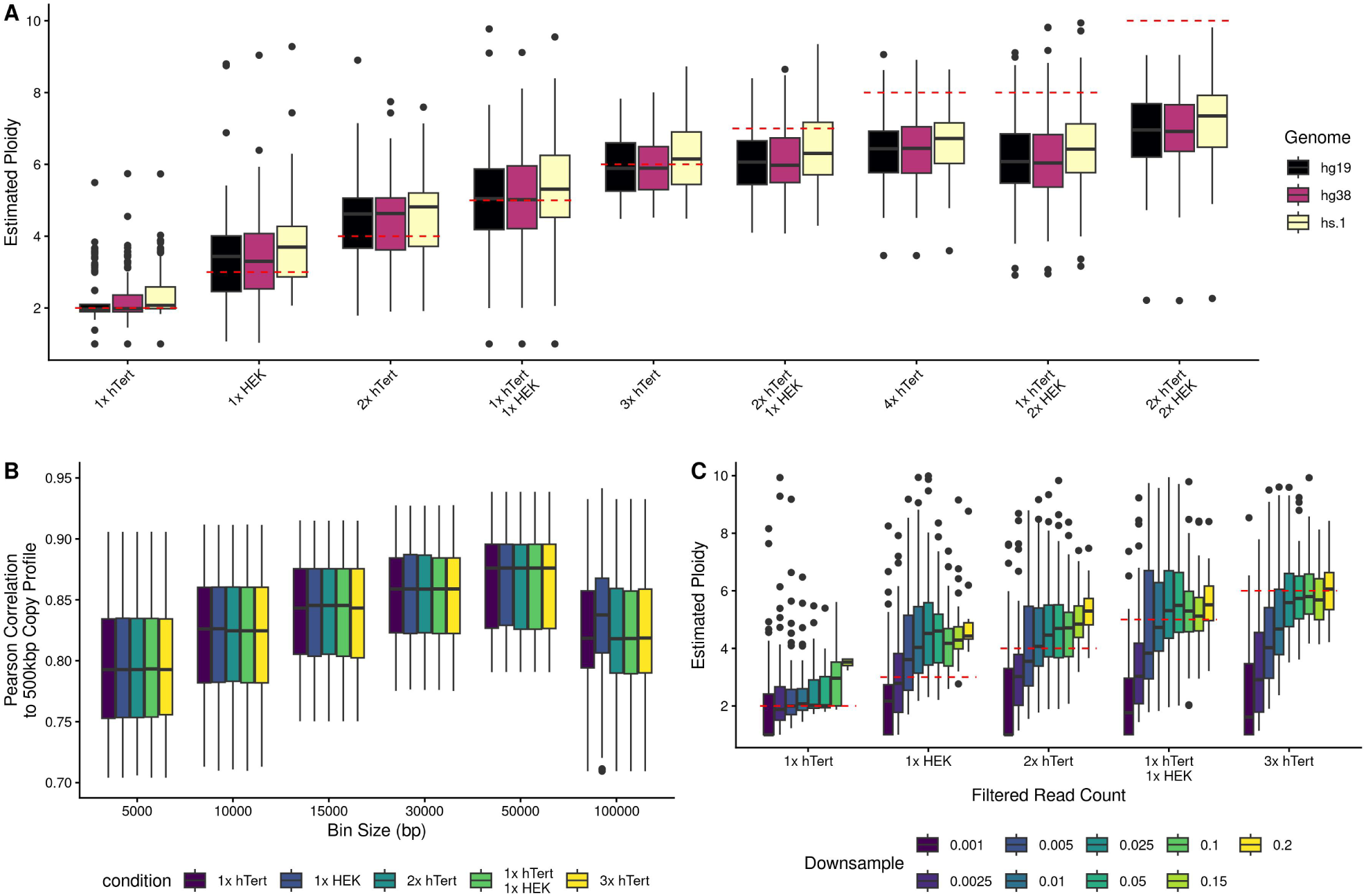
(A) Estimate Average Ploidy Across the Ploidy Ladder Comparing reads aligned to hg19, hg38, and hs.1. Dashed red line indicates expected ploidy based on the number of cells spotted in each well. (B) Pearson Correlation of the copy number profile compared to 500kbp bin sizes. (C) Estimated Average Ploidy across the ladder across different target depth. Higher target depths exclude cells which do not originally have the required depth.

**Supplementary Figure 4:**
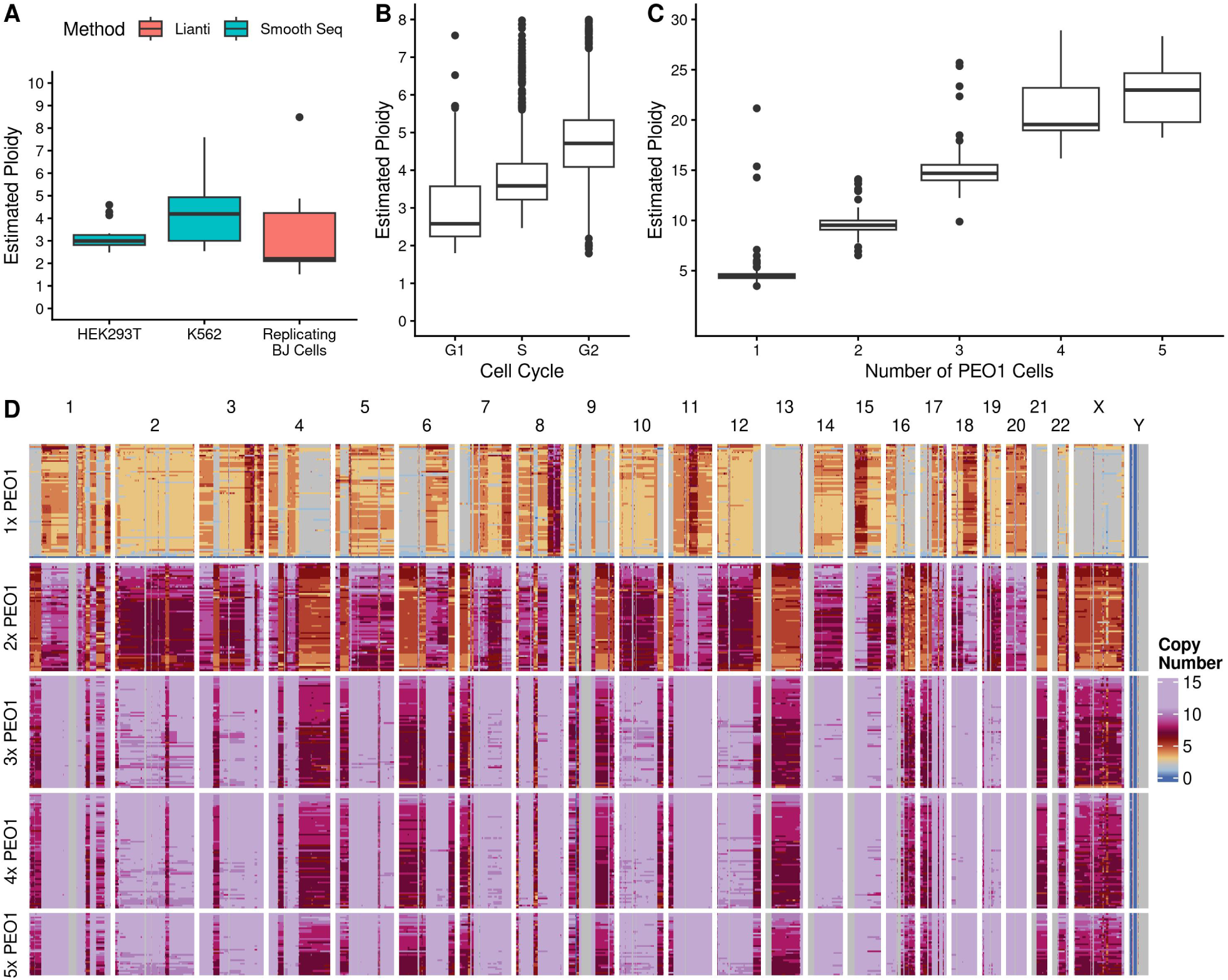
(A) Estimated Ploidy for cells sequenced using Smooth Seq and Lianti. (B) Estimated average ploidy for GM18507 cells spotted published in the original DLP+ paper [20]. (C, D) Estimated average ploidy and final copy number states across the PEO1 cell spotting ladder published in the scAbsolute paper [25].

